# DAVID: An open-source platform for real-time emotional speech transformation: With 25 applications in the behavioral sciences

**DOI:** 10.1101/038133

**Authors:** Laura Rachman, Marco Liuni, Pablo Arias, Andreas Lind, Petter Johansson, Lars Hall, Daniel Richardson, Katsumi Watanabe, Stéphanie Dubal, Jean-Julien Aucouturier

**Affiliations:** Science and Technology of Music and Sound STMS UMR 9912 / Institut de Recherche et Coordination en Acoustique et Musique, IRCAM, 75004 Paris, France; CNRS UMR 7225 / Sorbonne Universités, UPMC Univ Paris 06 UMR S 1127, Paris, France; Inserm U 1127, Paris, France; Institut du Cerveau et de la Moelle Épinière, ICM, F-75013, Paris, France; Lund University Cognitive Science, Lund University, 221 00 Lund, Sweden; Swedish Collegium for Advanced Study, Uppsala University, 752 38 Uppsala, Sweden; Department of Experimental Psychology, University College London, London WC1E 6BT, United Kingdom; Department of Intermedia Art and Science, Faculty of Science and Engineering, Waseda University, 169-8555 Tokyo, Japan; Research Center for Advanced Science and Technology, the University of Tokyo, 153 8904 Tokyo, Japan

## Abstract

We present an open-source software platform that transforms the emotions expressed by speech signals using audio effects like pitch shifting, inflection, vibrato, and filtering. The emotional transformations can be applied to any audio file, but can also run in real-time (with less than 20-millisecond latency), using live input from a microphone. We anticipate that this tool will be useful for the study of emotions in psychology and neuroscience, because it enables a high level of control over the acoustical and emotional content of experimental stimuli in a variety of laboratory situations, including real-time social situations. We present here results of a series of validation experiments showing that transformed emotions are recognized at above-chance levels in the French, English, Swedish and Japanese languages, with a naturalness comparable to natural speech. Then, we provide a list of twenty-five experimental ideas applying this new tool to important topics in the behavioral sciences.

## 1 Introduction

The use of well-defined stimulus material is an important requirement in experimental research, allowing for replicability and comparison with other studies. For this reason, researchers interested in the perception of emotions often use datasets of stimuli previously validated with affective norms. An increasing number of datasets exist for both facial expressions (e.g. the Karolinska Directed Emotional Faces - 70 individuals, each displaying 7 facial expressions, photographed from 5 different angles, Goeleven et al, 2008), vocal expression (e.g. the Montreal Affective Voices - 10 actors, each recording 9 non-verbal affect bursts, Belin et al, 2008) and musical extracts (e.g. The Montreal Musical Bursts - 80 short musical improvisations conveying happiness, sadness or fear, Paquette et al, 2013).

However, using datasets of static stimuli, regardless of how well controlled, comes with a number of generic limitations. First, such datasets leave researchers only little control over the para-emotional parameters of the expressions (e.g., the specific person expressing the emotion or the verbal content accompanying the expression). Some research questions may require more control over the stimulus material: for instance, to investigate social biases, one may want a certain emotion to be expressed by both a young girl and an older woman with the exact same acoustic cues. Second, actor-recorded stimuli do not allow for fine control over the intensity with which emotions are expressed: e.g., some actors may be more emotionally expressive than others, or perhaps more expressive when it comes to happiness than sadness. In an attempt to control for such parameters, various researchers have used morphing techniques between e.g. a neutral and an emotional facial expression (Sato et al, 2004), or between two different emotional vocal expressions (Bestelmeyer et al, 2012). Morphings can gradually increase the recognizability of an emotional stimulus or create arbitrarily ambiguous emotional voices (Bestelmeyer et al, 2012). However, morphings do not only affect expressive cues that are involved in the communication of emotion, but also cues that may not be linked directly to emotions, or that one may not want to be morphed. For instance, it is typically impossible to only morph the pitch, but not the loudness, between two emotional voices. Moreover, with morphings, the para-emotional context (e.g., specific speakers) remains limited to the stimuli that are included in the database. A final generic limitation of pre-recorded datasets is that they necessarily only consist of third-person stimuli. However, in many experimental contexts, one may desire to control the emotional expression of the participants themselves, and not that of unknown actors. For example, social psychology researchers may want to study participants’ interactive behavior while controlling whether they sound positive or negative. It remains difficult to create such situations without demand effect, e.g. not asking or otherwise leading participants to "act" happy or sad.

Rather than a data set of controlled emotional *stimuli*, it would therefore be useful to have a data set of controlled emotional *transformations*, that can be applied to arbitrary stimulus material while still preserving well-defined properties of recognizability, intensity and naturalness. Such data sets exist in the visual domain, for the synthesis of facial expressions. For instance, tools have been developed that can very precisely manipulate facial cues to alter perceived personality traits (Todorov et al, 2013) or the emotional expression (Roesch et al, 2011) of computer-generated or digitized faces, allowing for a high level of control. However, no such tools exist in the domain of vocal expression to the best of our knowledge. More precisely, while emotional voice synthesis is an active research field in the audio engineering community, no such tool comes with the experimental validation and technical requirements necessary for psychological research. Voice is a powerful medium for the expression of emotion (Bachorowski and Owren, 1995; Juslin et al, 2005). With a suitable voice transformation tool, it should be possible to change the emotional expression of speech after it is produced and, if computed fast enough, the transformations could even appear to occur in "real-time". With such a tool, one would be able to modify vocal emotional expressions in live and more realistic settings and study not only the perception of emotions in third party stimuli, but also the perception of self-produced emotions, opening up a vast amount of experimental questions and possibilities.

In this article, we present DAVID^1^, a novel open-source software platform providing a set of programmable emotional transformations that can be applied to vocal signals. The software makes use of standard digital audio effects, such as pitch shift and spectral filtering, carefully implemented to allow both realistic and unprecedentedly fast emotional transformations (section 2). DAVID was used in a previous study by Aucouturier et al (2016) where participants read a short text while hearing their voice modified in real-time to sound more happy, sad or afraid. Results of this study showed that a great majority of the participants did not detect the manipulation, proving that the emotional transformations sounded natural enough to be accepted as self-produced speech and that they were fast enough to allow for uninterrupted speech production. In addition, participants’ mood ratings changed in the same direction as the manipulation, suggesting that the transformations carry some emotional meaning.

Extending beyond this first experiment, we present here results from an additional series of experimental studies that validate four important claims that we consider indispensable for the tool to be useful in a larger set of psychological and neuroscience research paradigms, namely that the transformations are recognizable, natural, controllable in intensity, and reasonably intercultural (see section 3). Based on these results, we propose here a tentative list of 25 application ideas in a selection of research areas where we argue this new transformation software will be of particular importance.

## 2 Emotional transformations

### 2.1 Emotional speech synthesis techniques

Consciously or not, we convey emotional information with our speech. The words and syntactic structures that we use reveal our attitudes, both towards the topic of conversation and towards the person we converse with. Besides words, the sole *sound* of our voice is rich in information about our emotional states: higher fundamental frequency/pitch when happy than sad (Scherer and Oshinsky, 1977), faster speech rate when excited, raising intonation/prosody when surprised (Banziger and Scherer, 2006). Computerized audio analysis and synthesis are important techniques to investigate such acoustic correlates of emotional speech (Scherer, 2003). Widely-used phonetical analysis tools like Praat (Boersma and Weenink, 1996) allow the automatic analysis of large corpus of speech in terms of pitch, duration and spectral parameters (Laukka et al, 2005). More recently, speech synthesis techniques, typically pitch-synchronous overlap-and-add methods (PSOLA) and shape-invariant phase vocoder (Roebel, 2010), support the active testing of hypotheses by directly manipulating the acoustic parameters of the vocal stimuli (Bulut and Narayanan, 2008).

Beyond its use for psychological experimentation, emotional speech synthesis is now a widely researched technique per se, with applications ranging from more expressive text-to-speech (TTS) services for e.g. automatic ticket booths (Eide et al, 2004) to restoration of voices in old movies (Bertini et al, 2005) or more realistic non-player characters in video games (Farner et al, 2008). One major concern with such systems is the degree of realism of the synthesized voice. In early attempts, this constraint was simply relaxed by designing applications that did not need to sound *like anyone in particular* : for instance, cartoon baby voices for entertainment robots (Oudeyer, 2003). For more realism, recent approaches have increasingly relied on modifying pre-recorded units of speech, rather than synthesizing them from scratch (but see Astrinaki et al, 2012). One of such techniques, concatenative synthesis, automatically recombines large numbers of speech samples so that the resulting sequence matches a target sentence and the resulting sounds match the intended emotion. The emotional content of the concatenated sequence may come from the original speaking style of the pre-recorded samples ("select from the sad corpus") (Eide et al, 2004), result from the algorithmic transformation of neutral samples (Bulut et al, 2005), or from hybrid approaches that morph between different emotional samples (Boula de Mareüil et al, 2002). Another transformation approach to emotional speech synthesis is the recent trends of "voice conversion" research, which tries to impersonate a target voice by modifying a source voice. This is typically cast as a statistical learning task, where the mapping is learned over a corpus of examples, using e.g. Gaussian mixture models over a parameter space of spectral transformation (Inanoglu and Young, 2007; Godoy et al, 2009; Toda et al, 2012).

The tool we propose here, a voice transformation technique to color a spoken voice in an emotional direction which was not intended by the speaker, is in the direct tradition of these approaches, and shares with them the type of audio transformation used (i.e. temporal, pitch and spectral) and the need for high-level quality. However, we satisfy a very different constraint: the transformed voice has to be a realistic example of its speaker’s natural voice. Previous approaches have attempted-and succeeded - to produce either a realistic third-person voice (e.g. a considerate newscaster - Eide et al (2004)) or an exaggerated first-person (e.g. me as a happy child, me as an angry monster - Mayor et al (2009)). We describe here a technique which synthesizes a *realistic first-person* : me when I’m happy, me when I’m sad. We refer to the transformation as "natural", in that it effectively imparts the impression of a specific emotion for the listeners while being judged to be as plausible as other, non-modified recordings of the same speaker.

A second particularity of this work is that the transformation can be done in real-time, modifying speech as it is uttered, without imparting any delay capable of breaking a natural conversation flow (in practice, less than 20ms). This differentiates from previous work in several ways. First, the expectation of high realism has compelled previous approaches to design increasingly sophisticated analysis methods - time-domain PSOLA, Linear Prediction PSOLA (Moulines and Charpentier, 1990), Linear-Prediction Time-scaling (Cabral and Oliveira, 2005), Wide-band harmonic sinusoidal modeling (Mayor et al, 2009), to name but a few. As a consequence, none of these approaches can meet real-time constraints, especially as predictive models require a short-term accumulator of past data (but see Toda et al, 2012; Astrinaki et al, 2012, for recent progress on that issue). Second, many techniques rely on strategies that are incompatible with the real-time following of an input voice: speeding the voice up or down (Cabral and Oliveira, 2005), or inserting paralinguistic events such as hesitation markers <ERR> or <AHEM>. The approach described here manages to operate in real-time by careful design rather than by technical prowess. First, we favor effects that can be implemented efficiently, such as simple time-domain filtering, and in cascade (such as vibrato and pitch shifting both using the same pitch shifting module). Second, because the manipulation is designed to be "natural", our effects operate over very subtle parameter ranges (e.g. +/- 40 cents pitch shifting, instead of e.g. +/- 1 octave as targeted in Cabral and Oliveira, 2005), for which even simplistic (and fast) approaches are sufficient.

### 2.2 Software distribution

DAVID is a software platform developed to apply audio effects to the voice both online and offline. The platform provides four types of audio effects, or building blocks, that can be combined in different configurations to create several emotions: happy, sad, and afraid (and more are possible). DAVID is implemented as an open-source patch in the Max environment (Cycling74), a programming software developed for music and multimedia. The DAVID software and accompanying documentation can be downloaded under the MIT license from http://cream.ircam.fr. Using DAVID first requires to install the Max environment, which is provided in free versions for Windows and Mac systems. DAVID comes with the parameter presets used in the validation studies described below, but users also have full control over the values of each audio effect to create their own transformations and store them as further pre-sets. The present article is based on software version v1.0 of DAVID (release date: 15/10/2015), see the DAVID website for further updates and new functionalities.

### 2.3 Algorithms used in DAVID

DAVID is designed as a collection of building blocks, or "audio effects", that can be combined in different configurations to create emotion transformations. Each audio effect corresponds to a frequently-identified correlate of emotional voices in the literature. For instance, fear is often associated with fluctuations in the voice pitch (Laukka et al, 2005) - an effect we implement here as vibrato (see below). However, we choose not to associate an individual effect with a individual emotion (e.g. vibrato ↔ fear), because we observed a large degree of overlap and/or contradicting claims in previous works. For instance, Laukka et al (2005) observe that "a low mean pitch is a correlate of positive valence, but also of negative arousal", casting doubt on what should be associated with a state of joy. Rather, audio effects in DAVID are best described as "things that often happen to one’s voice when in an emotional situation". How these effects map to emotions depends on the way the effects are quantified, the way emotions are represented (words, multidimensional scales, etc.), and possibly other factors such as context or culture (Elfenbein and Ambady, 2002), and elucidating this mapping is not the primary concern of our work.

In the experiments presented here, we tested three types of transformations - happy, sad and afraid - each composed of several, sometimes overlapping audio effects (e.g. afraid and happy both include the inflection effect). The audio effects used in each manipulation are listed in Table 1, and their algorithmic details given below.

**Table 1.** List of the atomic digital audio effects used in this work, and how they are combined to form emotional transformations happy, sad and afraid.

| Effects |  | Transformations |  |  |
| --- | --- | --- | --- | --- |
|  |  | Happy | Sad | Afraid |
| Time-varying | Vibrato |  |  | ✓ |
|  | Inflection | ✓ |  | ✓ |
| Pitch shift | Up | ✓ |  |  |
|  | Down |  | ✓ |  |
| Filter | High-shelf<br>(“brighter”) | ✓ |  |  |
|  | Low-shelf<br>(“darker”) |  | ✓ |  |

In addition to being consensual emotional correlates, the audio effects described below obey several constraints to make them usable in real-time. First, none of them can alter the speed of speech delivery. Varying speech speed is a commonly observed correlate of emotional voices, e.g. sad voices tend to be slower and happy voices faster (Scherer and Oshinsky, 1977). However, playing speech faster in real-time is impossible by construction (we cannot playback something which has not been recorded yet); playing it slower is technically feasible using a buffer memory, but will result in de-synchronization with the speaker and will immediately be noticeable after a few tens of milliseconds of delay (let alone after several minutes of speech) (see e.g. Badian et al, 1979).

Second, none of the audio effects can rely on contextual knowledge, such as linguistic units and boundaries.

For instance, happy voice prosody tend to raise in pitch at the end of sentences (Hammerschmidt and Jurgens, 2007). However, detecting this in real-time requires to process larger segments of audio, to observe a decrease in the pitch contour, with a consequent augmentation of the system’s latency.

Third, all effects must have a computational complexity suitable for real-time. In addition to the latency due to the hardware configuration, each audio effect in the signal data flow give rise to further delay, which depends on the effect’s algorithmic complexity and the processor speed. Effects acting on individual audio samples (such as applying a square root function to the signal) are fastest; effects which require a buffered transform, such as FFT, are longest. We will give such implementation details in section 2.3.5. In all of our experiments, and for all manipulations, we reached global latencies below 20 ms.

#### 2.3.1 Pitch shift

Pitch-shift denotes the multiplication of the pitch of the original voice signal by a constant factor α. Increased pitch (α > 1) often correlates with highly aroused states such as happiness, while decreased pitch (α < 1) correlates with low valence, such as sadness (Laukka et al, 2005).

##### Implementation

In DAVID, pitch-shift is implemented as a standard application of the harmonizer, i.e. a time-varying delay. For this, a maximum delay time has to be specified, that consequently defines the needed amount of memory in order to delay the incoming signal, and thus the latency of the algorithm (this parameter is accessible as *window* in DAVID). Pitch is shifted by a constant factor (see figure 1a). In order to reduce computational load, early processing stages of the constant pitch-shift algorithm are shared with the time-varying vibrato and inflection, and factors for multiplying pitch are accumulated where appropriate.

**Figure 1.**
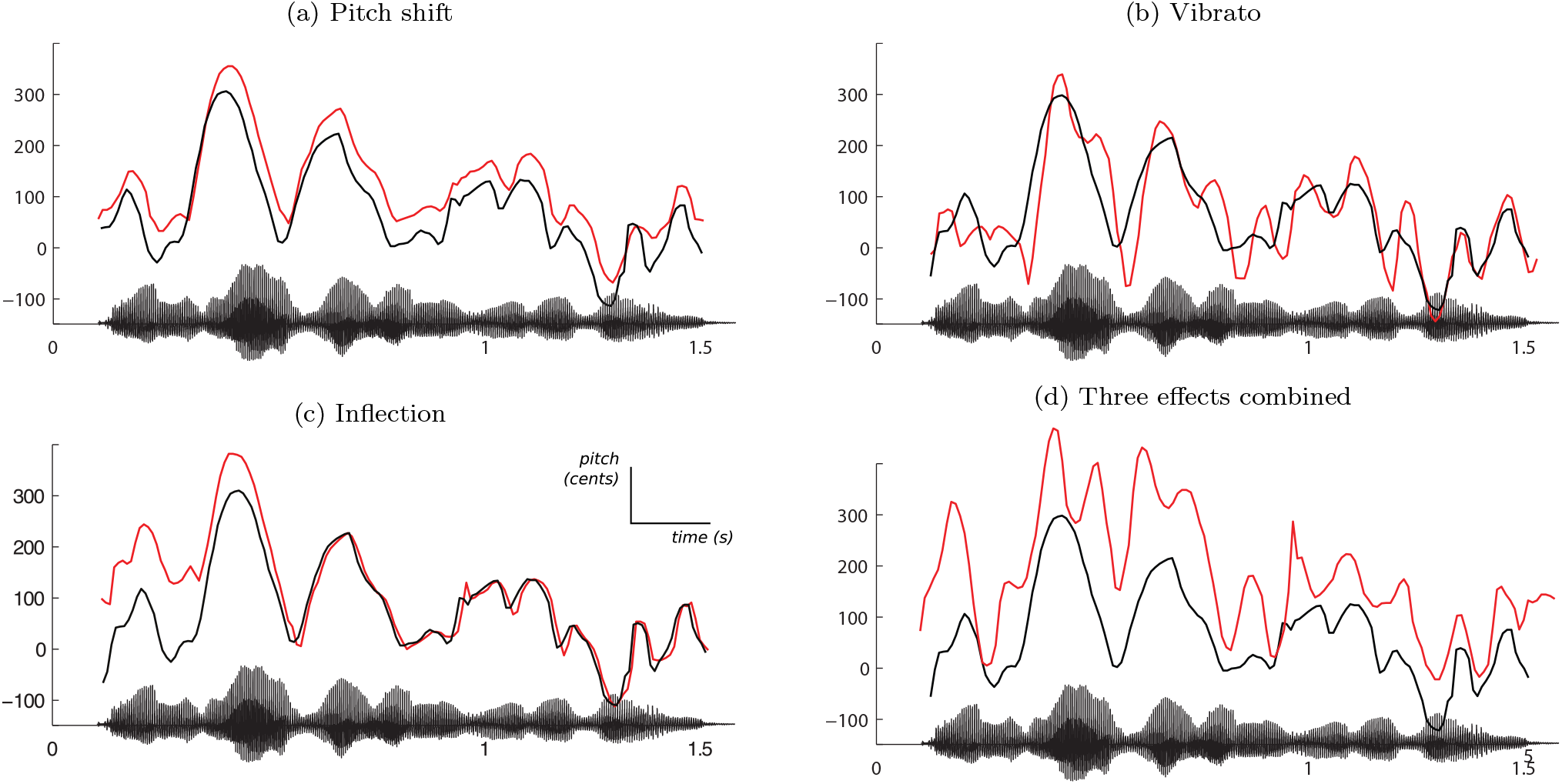
Three of the audio effects available in DAVID, applied on the same recording by a French female speaker, saying “*Je suis en route pour la réunion*” (I’m on my way to the meeting). The solid black line represents the time series of pitch values in the original recording (estimated with the SWIPE algorithm - Camacho and Harris (2008)) and the red line represents the pitch of manipulated audio output. The speech waveform of the unmodified recording is shown on the x-axis of each subfigure. Pitch values on y-axis are normalized to cents with respect to mean frequency 200Hz. (a) The pitch is shifted upwards by 40 cents. (b) Vibrato is applied with a rate of 8.5 Hz and a depth of 40 cents. (c) Inflection kicks in at the start of the utterance, with an initial shift of +140 cents, and recedes after 500 ms (implemented in the happy transformation). (d) The three effects combined, for illustration purposes. The audio effects can be applied in any configuration.

##### Parameters

Pitch-shift is used in the happy transformation with a positive shift of + 50 cents (i.e., one half of a semitone), and in the sad transformation with a negative shift of-70 cents. The maximum delay time is set by default to 10 milliseconds

#### 2.3.2 Vibrato

Vibrato is a periodic modulation of the pitch (fundamental frequency) of the voice, occurring with a given rate and depth. Vibrato, also related to jitter, is frequently reported as a correlate of high arousal (Laukka et al, 2005).

##### Implementation details

Vibrato is implemented as a sinusoidal modulation of the pitch shift effect, with a rate parameter (modulation frequency, in Hz), a depth (in cents) and a random variation of the rate (in percentage of the rate frequency). Figure 1b shows a typical output of the algorithm (using a speech extract from our experimental data).

##### Parameters

The afraid transformation uses vibrato with a rate of 8.5Hz, a depth of +/- 40 cents and a 30% random rate variation.

#### 2.3.3 Inflection

Inflection is a rapid modification (∽500ms) of the pitch at the start of each utterance, which overshoots its target by several semitones but quickly decays to the normal value. Inflection is associated with several speech disorders and evocative of stress and negative valence (Scherer, 1987).

##### Implementation details

DAVID analyses the incoming audio to extract its root-mean-square (RMS), using a sliding window. When the RMS reaches a minimum threshold, the system registers an attack, and starts modulating the pitch of each successive frame with a given inflection profile (see figure 1c). The inflection profile can be specified by the user, together with a minimum and maximum pitch shift, as well as a duration.

##### Parameters

Two inflection profiles are proposed: the first, associated in our experiments to the happy transformation, quickly increases from-200 cents to 140 cents, then decaying to the original pitch over a total duration of 500 ms; the second, associated to the afraid effect, is a sinusoidal curve between-200 and 200 cents with a duration of 500 ms, starting at its maximum position and decaying to the original pitch.

#### 2.3.4 Filtering

Filtering denotes the process of emphasizing or attenuating the energy contributions of certain areas of the frequency spectrum. The acoustics of emotional speech are rarely analyzed in terms of global spectral changes (Pittam et al, 1990), however, we found that some simple filtering is often successful in simulating behaviors of vocal production that are typically associated with emotional content. For instance, high arousal emotions tends to be associated with increased high frequency energy, making the voice sound sharper and brighter (Pittam et al, 1990); this can be simulated with a high-shelf filter. Conversely, "sad" speech is often described as darker, a perception simulated with a low-shelf filter.

##### Implementation details

All filters are implemented as 5-order Butterworth IIR filters. Filter design is done offline (not in real-time), with a bilinear transform.

##### Parameters

The happy transformation uses a high-shelf filter with a cut-off frequency at 8000 Hz, +9.5 dB per octave ("brighter"). The sad transformation uses a low-shelf filter with a cut-off frequency at 8000 Hz,-12 dB per octave ("darker").

### 2.3.5 System and algorithm latency

Figure 2 gives a schematic explanation of the latencies involved in the realization of our real-time audio processing system. The incoming audio has to be captured and converted from analog to digital format before reaching the memory of the application. This causes a first delay (input Δ_*t*_). Similarly, after all processing is done, the digital signal has to be routed back from application memory to the output device, undergoing digital to analog conversion - hence an output Δ_*t*_. Both input and output delays (also known as *roundtrip* latency) occur even if no processing is done: this is the delay time that is typically experienced when talking into a microphone plugged into the sound card, while listening back to the voice through headphones. Roundtrip latency depends on the system s hardware and software, and can be easily optimized to the range 2-5ms MacMillan et al (2001). However, when some processing is applied, a low latency can affect the output sound quality, because the high rate at which computer and audio card exchange data may provide less samples than needed for some algorithms to achieve a correct result. In the Max environment, the exchange rate between the CPU and the sound card is controlled by means of the *I/O vector size* (which corresponds to the input and output Δ_*t*_), while the *signal vector size* determines the exchange rate within Max itself.

**Figure 2.**
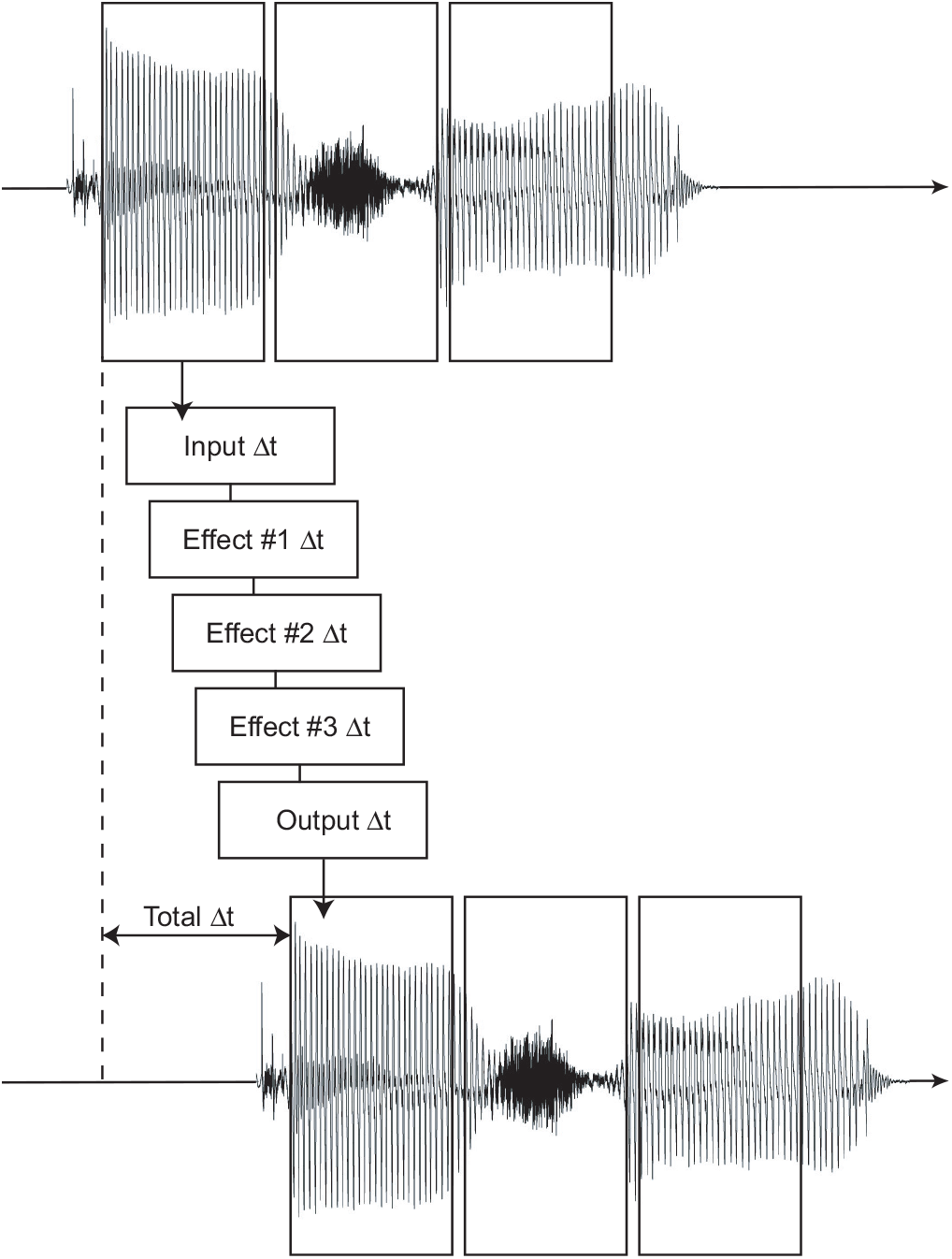
Illustration of the delays involved in the realization of our real-time audio processing system. Beyond a baseline I/O latency (input and output Δ_*t*_), each atomic effect in the signal data flow (3 as illustrated here) imparts further delay, which depends on the effect’s algorithmic complexity.

Our recommended set-up for using DAVID in a realtime context consists of:

- a high-end computer, such as 2,5 GHz Intel Core i7 MacBook Pro, running Mac OS X 10.10.2
- a medium to high-end external audio interface, such as a RME UCX Fireface sound card
- I/O vector size: 256 samples
- Signal vector size: 128 samples

Using this configuration, we consistently measured a roundtrip latency of 9.5 ms.

As for the algorithmic latency, all of the modules in DAVID are based on the same pitch shifting engine, that is the harmonizer described in section 2.3.1. The only latency is thus given by the maximum delay time in the harmonizer, that is set by default to 10 milliseconds. Adding this latency to the roundtrip one, the global default latency provided by DAVID amounts to 19.5 milliseconds.

## 3 Experimental validation of claims 1-4

We present here results from a series of experimental studies that validate four important claims that we consider indispensable for the tool to be useful in psychological and neuroscience research:

**Claim 1:** The emotional tone of the transformed voice is recognizable.
**Claim 2:** The transformed voices sound natural and are not perceived as synthetic.
**Claim 3:** The emotional intensity of the transformed voices can be controlled.
**Claim 4:** The emotional transformations can be applied in several languages, making the tool applicable in multiple research environments, as well as to cross-cultural research questions.

### 3.1 Stimuli

We recorded six neutral sentences spoken by 12 French (6 female), 9 English (4 female), 14 Swedish (7 female), and 11 Japanese (7 female) speakers. The sentences were chosen from a database that included semantically neutral sentences (Russ et al, 2008). Speakers were asked to produce each sentence eight times with a neutral expression and three times with each of the emotional expressions (happy, sad, afraid). The recordings took place in a sound-attenuated booth, using GarageBand software (Apple Inc.) and an omnidirectional headset microphone (DPA d:fine 4066) connected to an Apple Macintosh computer. Digitization of the recordings was done at a 44.1 kHz sampling rate and 24-bit resolution. Based on the quality of the recordings, six speakers (three female) and four sentences were selected in each language. This selection was finally included in three behavioral experiments to test the validity of the software tool, yielding 24 different speaker-sentence combinations per language.

For each speaker and sentence, we selected the first four neutral recordings for further processing. If the quality was insufficient, we selected the next available recording. For the sentences spoken with an emotional tone, we selected only one recording.

Three out of the four neutral recordings were processed with our tool to transform the voices into happy, sad, and afraid voices. For each emotion, we selected the parameters for the audio effects such that we judged the emotional transformation to be recognizable, yet natural. In the remainder of this article, we will refer to these parameter settings, indicated in section 2 and in Table 2, as the "nominal level" Furthermore, we processed the recordings with the same audio effects at two increasingly reduced intensity levels. We thus tested three emotional transformations at three intensity levels. All audio files were normalized for maximum peak intensity using Audacity version 2.1.0.

**Table 2.**
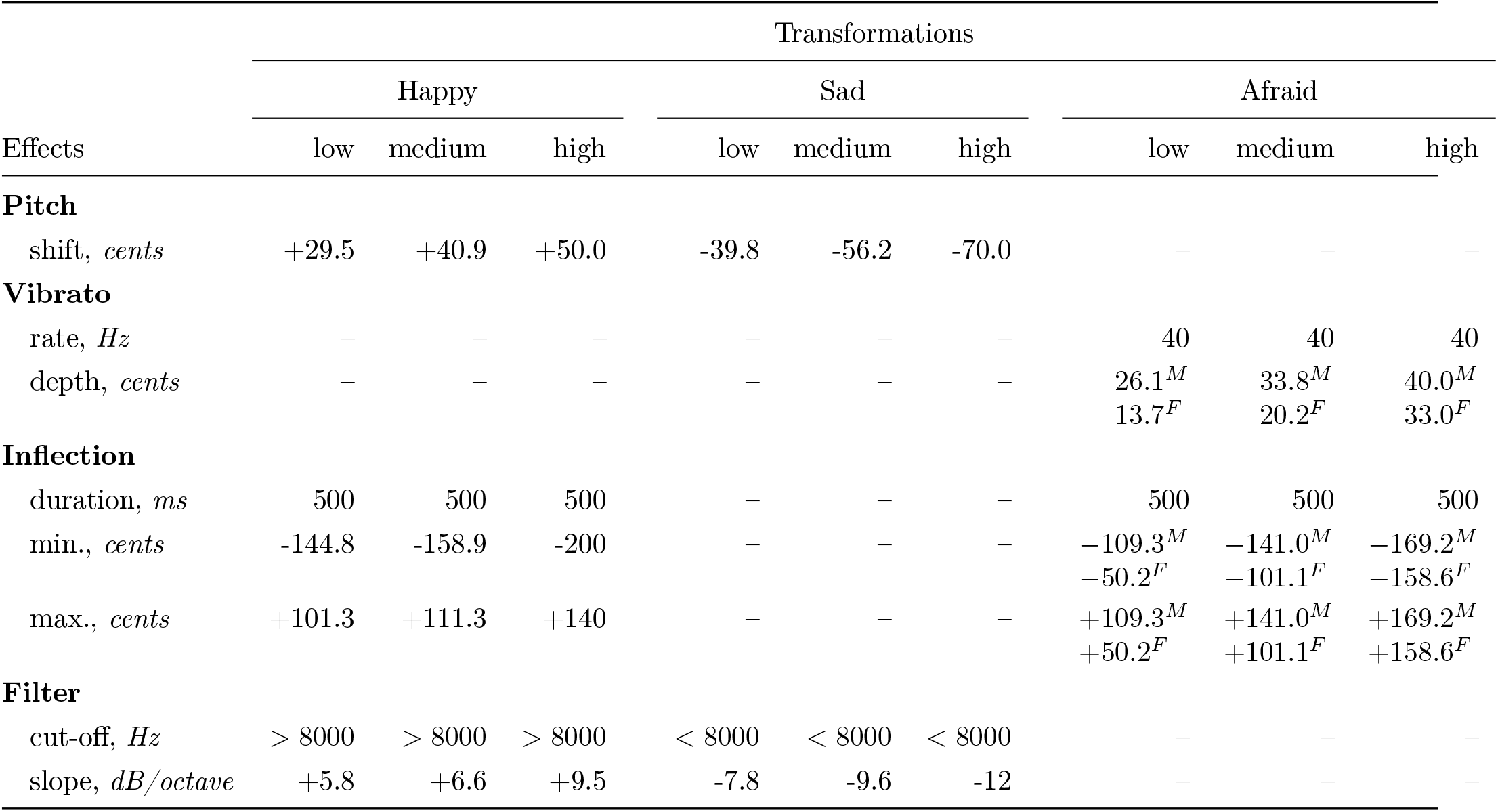
List of the parameters used in the validation experiments. For the afraid transformation different values were used for male and female voices, due to strong differences of the audio effects depending on the gender of the speaker.

### 3.2. Methods

#### 3.2.1 Participants

The validation studies of the emotional voice effects were carried out in four languages: French, English, Swedish and Japanese, in IRCAM (France), University College London (UK), Lund University (Sweden) and Waseda University (Japan). Participants in the study comprised 20 native French volunteers (mean age = 25.4 years, SD = 4.9, 10 females), 27 native British English volunteers (mean age = 26.1 years, SD = 5.6, 17 females), 20 native Swedish volunteers (mean age = 28.1 years, SD = 5.3, 10 females), and 20 native Japanese volunteers (mean age = 21.1 years, SD = 1.4, 10 females). Two female English participants were excluded because they did not satisfy age or language requirements. Furthermore, responses of one female English volunteer were not recorded during the emotion recognition task (see below) due to technical problems. Volunteers were recruited through local databases and mailing lists in the respective countries and were financially reimbursed for their participation. The study was approved globally by the IRB of the French Institute of Health and Medical Research (INSERM), as well as locally by the departmental review boards of University College London, Lund University and Waseda University. All participants gave written informed content.

### 3.3 Procedure

To validate Claims 1-4, participants performed three consecutive tasks: A naturalness rating task (Claim 2), an emotion recognition task (Claim 1), and an intensity rating task (Claim 3). Participants always performed all three tasks in the aforementioned order to avoid an interference effect of the recognition of the emotional transformations on naturalness ratings. We ran these validation experiments in the three different languages to address Claim 4. Together, the three experiments took approximately one hour. The voice stimuli were presented through closed headphones (Beyerdynamics, DT770, 250 ohm), with the sound level adjusted by the participant before the start of the experiment. An Apple MacBook Pro running PsychoPy (Peirce, 2007) was used to control stimulus presentation and the recording of responses.

#### Emotion recognition task

In each trial, participants listened to two utterances of the same sentence and the same speaker. The first utterance was always the neutral recording and the second utterance was either the same recording unprocessed (neutral condition), or processed with one of the emotional transformations. We only used the neutral recordings in this experiment, the naturally produced emotions were used in the other two tasks described below. Participants compared the two utterances in order to indicate in a forced choice task whether the second extract sounded happy, sad, afraid, or neutral. Participants could choose "none of the above" whenever they heard a difference that did not fit one of the response labels (e.g. because the voice did not sound emotional at all, or because it sounded more like another emotion or a mixture of different emotions). Participants could listen to the voices as many times as necessary to make their judgment before proceeding to the next trial.

#### Naturalness task

In this task, participants heard one voice, either natural or modified, per trial and rated the naturalness of the voice on a continuous scale from "not at all artificial/very natural" to "very artifical/not at all natural" (1-100). Participants were told that some of the voices were completely natural and that others were manipulated by a computer algorithm. As in the decoding task, participants could listen to each utterance as many times as needed to make their judgment.

#### Intensity task

In this task, like in the naturalness task, participants heard either a modified or a natural voice. In each trial the correct emotion label was presented on the screen and participants judged the emotional intensity on a continuous rating scale (1-100) from "not at all happy/sad/afraid" to "very happy/sad/afraid". In addition, participants rated the loudness (subjective sound intensity) of the utterance to avoid confusion between the emotional intensity and other perceived acoustic characteristics that are not necessarily related to the intensity of the emotion. Loudness ratings were not further analyzed.

### 3.4 Data analysis

We calculated the mean ratings of naturalness and intensity for the naturalness and intensity tasks. In addition, we computed mean accuracy scores for the emotion recognition task. To take a possible response bias in the recognition task into account, we calculated the unbiased hit rates *(H_u_)* and chance proportions (*p_c_*) for each participant (Wagner, 1993). We then compared the arc-sine transformed scores by means of paired *t* tests. For easier comparison with other studies, we also calculated the proportion index (pi) that transforms the recognition rates to a standard scale where a score of 0.5 equals chance performance and a score of 1.0 represents a decoding accuracy of 100 percent (Rosenthal and Rubin, 1989).

## 4 Results

We first tested claims 1-3 in the French population. These results are presented in sections 4.1, 4.2 and 4.3. We then present the results from the English and Swedish populations in section 4.4.

### 4.1 Claim 1: Emotional transformations can be recognized

Proportion indices of the raw accuracy scores (see figure 3a) fell within the range of accuracy rates reported in a review on vocal expression by Juslin and Laukka (2003) (this study, Juslin and Laukka; happy = 0.76, 0.51-1.0; sad = 0.83, 0.80-1.0; afraid = 0.70, 0.65-1.0). Paired *t* tests of the unbiased hit rates at the nominal level against the individual chance proportions showed that all three emotional effects were correctly recognized at rates above the individual chance level: happy, *t*(19) = 6.2; sad, *t*(19) = 5.5; afraid, *t*(19) = 5.6 (all significant after Holm-Bonferroni correction, *ps <* .001, see also table 3). Additionally, patterns of errors showed that both the happy and sad responses were more often chosen for the correct corresponding emotion than for any other emotion. However, while the afraid transformation was correctly recognized above chance level, it was also perceived as sadness in as many cases (Fig. 3b).

**Table 3.**
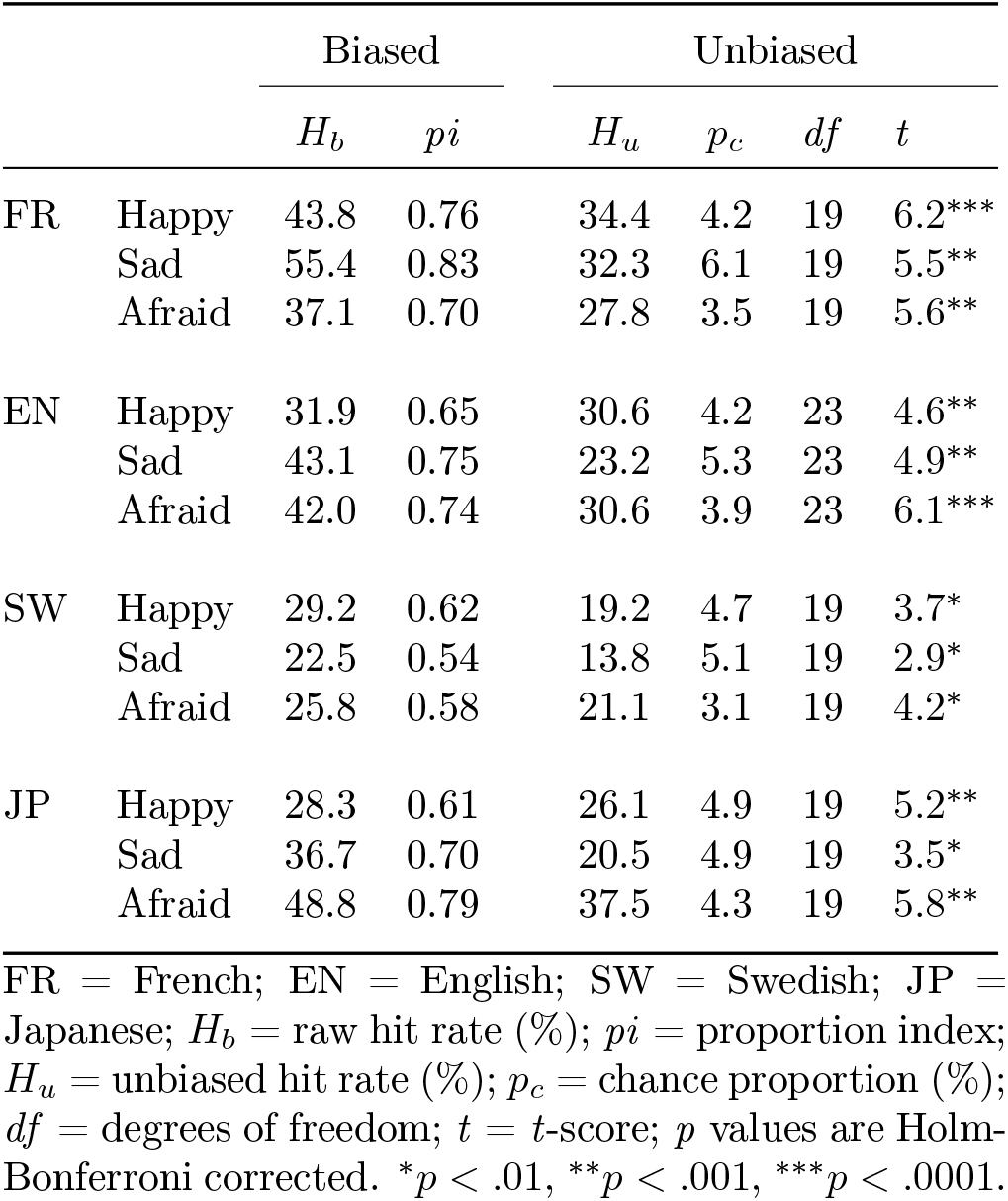
Emotion recognition scores, four languages

|  |  | Biased |  | Unbiased |  |  |  |
| --- | --- | --- | --- | --- | --- | --- | --- |
| | | $H_b$ | $pi$ | $H_u$ | $p_c$ | $df$ | $t$ |
| FR | Happy | 43.8 | 0.76 | 34.4 | 4.2 | 19 | 6.2*** |
|  | Sad | 55.4 | 0.83 | 32.3 | 6.1 | 19 | 5.5** |
|  | Afraid | 37.1 | 0.70 | 27.8 | 3.5 | 19 | 5.6** |
| EN | Happy | 31.9 | 0.65 | 30.6 | 4.2 | 23 | 4.6** |
|  | Sad | 43.1 | 0.75 | 23.2 | 5.3 | 23 | 4.9** |
|  | Afraid | 42.0 | 0.74 | 30.6 | 3.9 | 23 | 6.1*** |
| SW | Happy | 29.2 | 0.62 | 19.2 | 4.7 | 19 | 3.7* |
|  | Sad | 22.5 | 0.54 | 13.8 | 5.1 | 19 | 2.9* |
|  | Afraid | 25.8 | 0.58 | 21.1 | 3.1 | 19 | 4.2* |
| JP | Happy | 28.3 | 0.61 | 26.1 | 4.9 | 19 | 5.2** |
|  | Sad | 36.7 | 0.70 | 20.5 | 4.9 | 19 | 3.5* |
|  | Afraid | 48.8 | 0.79 | 37.5 | 4.3 | 19 | 5.8** |

FR = French; EN = English; SW = Swedish; JP = Japanese; $H_b$ = raw hit rate (%); $pi$ = proportion index; $H_u$ = unbiased hit rate (%); $p_c$ = chance proportion (%); $df$ = degrees of freedom; $t$ = $t$ -score; $p$ values are Holm-Bonferroni corrected. \* $p < .01$ , \*\* $p < .001$ , \*\*\* $p < .0001$ .

**Figure 3.**
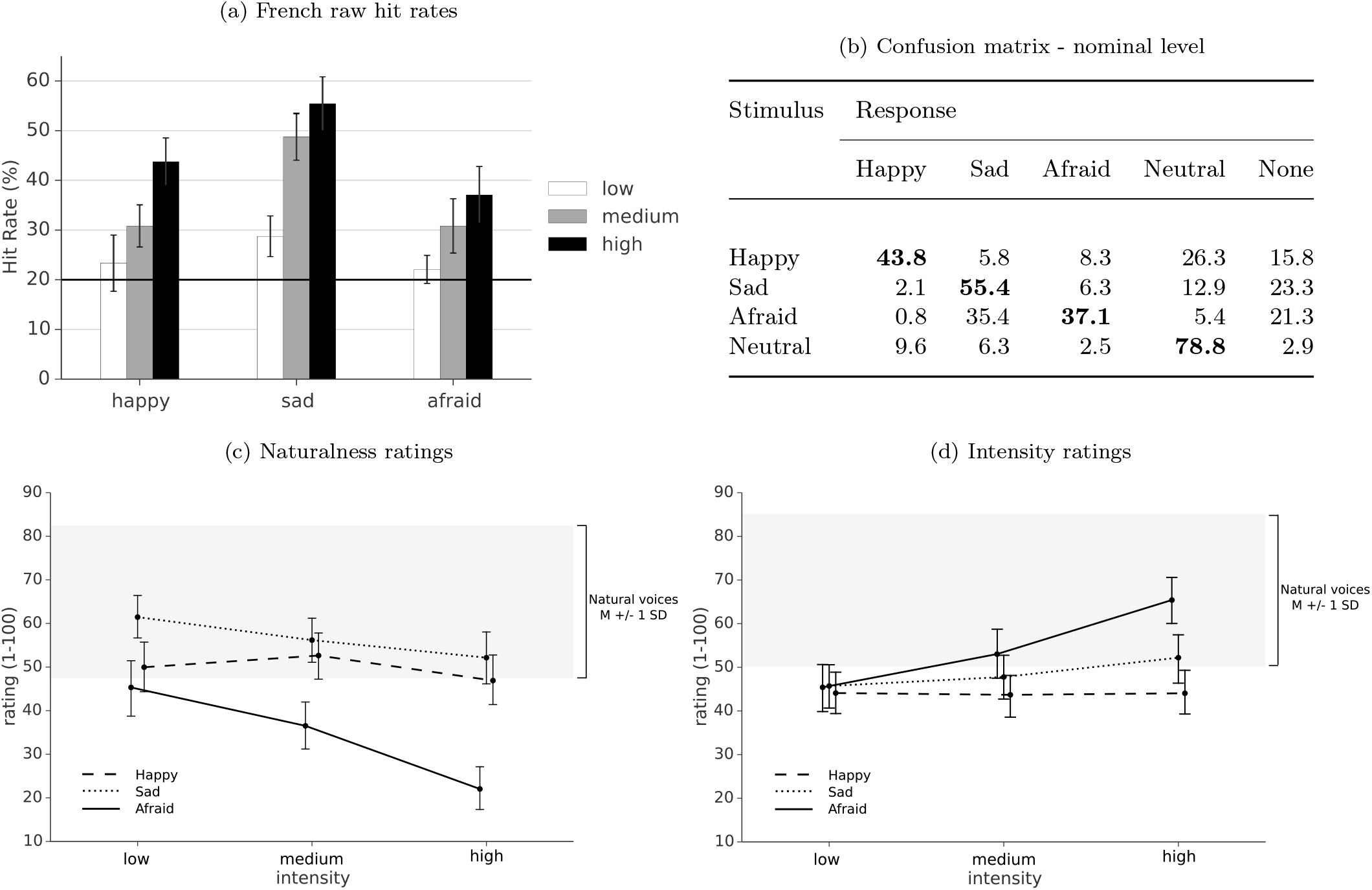
French results. (a) Raw accuracy scores for three emotions at the nominal level (’high’) and two lower intensity levels, error bars represent SEM, black line represents chance level (20%). (b) Confusion matrix shows the distribution (percentages) of responses at the nominal level. (c) Naturalness and (d) intensity ratings for three emotions at three intensity levels compared to unmodified voices (grey: mean ± 1 SD), error bars represent 95% confidence intervals.

### 4.2 Claim 2: Transformed voices sound natural

At the nominal level, the mean natural ratings were 46.9 for happy, 52.2 for sad, and 22.0 for afraid, with 95% confidence intervals [39.5, 54.3], [46.5, 57.9], and [15.2, 28.8], respectively (Fig. 3c). The mean naturalness rating of the sad transformation fell within one standard deviation of the mean naturalness rating for the naturally produced emotions (M = 64.9, SD = 17.5) (Fig. 3c), while the mean rating for happy fell just outside of this range. The afraid effect was rated as less natural (mean = 22.0). However, at decreasing intensity levels, it approached the naturalness range of naturally produced emotions (see also claim 3).

### 4.3 Claim 3: Control over emotional intensity

The intensity ratings of the afraid transformation showed a significant difference between the medium and highest levels, *t*(19) = 4.74, *p* = .0001 (Fig. 3d). There was also a significant difference between the lowest and medium levels,*t*(19) = 2.14, *p <* .05. The sad transformation showed a marginal statistical difference between the medium and highest levels, *t*(19) = 1.92, *p* = .07. The difference between the lowest and medium levels was not significant, low-medium, *t*(19) = -0.84, *p* = .41. There were no significant differences between the different intensity levels for the happy transformation: low-medium, *t*(19) = 0.20, *p* = .85; medium-high, *t*(19) = -0.12, *p* = .90.

### 4.4 Claim 4: Claims 1-3 hold in several languages

At the nominal level, unbiased hit rates were higher than chance level for all three emotions in French, English, Swedish and Japanese after Holm-Bonferroni correction (all *p*s<.01, see table 3). A two-way ANOVA with emotion as within-subject variable and language as between-subject variable showed a main effect of language, *F*(3,80)=2.869, *p* < .05. Fisher’s LSD post-hoc tests showed that this effect was driven by the Swedish participants who scored lower than all other samples. There was also a main effect of emotion, *F*(2,160)=3.631, *p* < . 05. Fisher’s LSD post-hoc tests revealed that the unbiased hit rates for the afraid transformation where higher than for both the happy and sad transformations. There was no significant interaction effect between language and emotion, *F*(6,160)=1.799, *p* = .10

Furthermore, French, English and Swedish data showed a similar pattern of naturalness ratings. Both the happy and sad effects were comparable to natural voices at the three levels whereas the naturalness of the afraid effect decreased at the two highest levels (see supplemental material). The Japanese data showed a similar pattern, although overall the emotional transformations were rated as less natural than the natural expressions.

Finally, intensity ratings showed similar patterns in all languages where ratings of both the afraid and sad effects increased at higher intensity levels and ratings for the happy effect remained the same for increasing intensity levels.

### 4.5 Incidental findings

Besides validating claims 1-4, the present study also allows for a number of incidental observations, which we discuss here in a post-hoc manner.

First, by varying the intensity of the effect from high to low, we found that the emotion recognition accuracy decreased, in all four languages. While this is not surprising, we are not aware of much prior literature on that issue, probably because of the difficulty to manipulate expression intensity in a systematic manner with actor-recorded stimuli.

Second, we found that intensity ratings for the happy voices did not increase with effect intensity, contrary to all other emotions. The finding held across all languages, and was especially intriguing given the gradual improvement in recognition rates when the transformation parameters increase. It seems that, while the happy effect is not perceived as more intense, it is more clearly perceived as happy. These results warrant further investigation of the respective contribution of different acoustical characteristics to emotional expressions.

Third, while the emotions were recognized above chance levels in all languages, we did find some differences in performance across languages. French proportion indices approach those reported in a meta analysis by Juslin and Laukka (2003), whereas the English, Swedish and Japanese data show overall slightly lower scores. In particular, recognition rates in the Swedish population were statistically lower than those in the three other populations. Based on the error patterns shown in the confusion matrices it seems possible that these differences can be attributed to the emotion labels and definitions used in the studies, of which the boundaries may have differed across languages. Swedish participants mistook an afraid voice for a sad one, and vice versa, more often in comparison to participants from the other populations. These discrepancies between different languages may therefore be a result of cultural differences, rather than perceptual differences.

## 5 Twenty-five applications in the behavioral sciences

We presented here a new software tool that is able to transform a neutral voice into either a happy, sad, or afraid voice. The transformations utilize well-known digital audio effects such as vibrato and spectral filtering. Furthermore, the transformations operate with a latency smaller than 20 ms and can thus be used in real-time contexts, for instance live interactions. In three consecutive listening experiments, we validated the following four claims. The emotional transformations were well recognized (Claim 1) and sounded as natural as non-modified expressions of the same speaker (Claim 2); the emotional intensity of the transformation could be controlled (Claim 3), and the transformations appeared valid in several languages (Claim 4), namely in French, English, Swedish and Japanese.

The main innovation of our approach, differentiating it from previous work, is that the manipulations are *both* real-time and natural. As already mentioned, previous attempts at real-time voice modification typically aimed to create caricatural or gender changes. Conversely, naturalness has been typically achieved so far in an offline context (e.g., pre-recorded samples), and has rarely been formulated in the context of voice change (a natural first person), but rather that of synthesis (a natural third person). Here, we achieve both simultaneously. This opens up new possibilities for experimental paradigms, from controlling the emotional vocal expression of one participant to studying the effect of emotional prosody in group interactions. We list a selection of these applications below.

### 5.1 Emotional prosody

1. Parametric variations: Traditional studies of emotional prosody typically rely on actor-produced impersonations of various emotions. While these constitute reliable stimuli to test listeners’ perception, they constitute a biased basis for the study of production, unlikely to span the whole register of possible prosodic variation. Using DAVID, one can parametrically sample a large, generic space of emotional prosodic variations (e.g., all vibrato frequency between 1Hz and 10Hz) and thus generate more ecologically valid templates of emotional expression, using e.g. reverse correlation techniques as employed with emotional faces (Mangini and Biederman, 2004).
2. Emotional intensity: Human actors have notorious difficulty manipulating the intensity of their emotional expression without a confounding variation of acoustic intensity or pitch (Ethofer et al, 2006). Consequently, the psychoacoustics of emotional intensity (what makes a happy voice happier) is still unknown to a large degree. Using DAVID, one can selectively manipulate the acoustical depth of various parameters (e.g. pitch shift independently from RMS energy), and examine how these parameter changes influence perceived emotional intensity.
3. Normalized task performance: Many psychophysical procedures require that participant performance be normalized at a standard level (typically d’ = .75 in 2AFC tasks) before other measures can be processed, e.g. to accurately estimate inflection points in psychometric curves, to generate templates of variation around a nominal point in reverse correlation methods, or to measure meta-cognitive capacity (Fleming and Lau, 2014). By systematically varying acoustic parameter strength, DAVID can be used to produce a continuous range of stimulus difficulties, e.g. in iterative procedures, and thus enable precise normalization of task performance regardless of individual differences.
4. Sensory vs emotional accuracy: Most studies investigating individual differences in emotional discrimination abilities use actor-produced stimuli. Such stimuli inherently confound individuals’ sensory performance (how well is a given individual able to perceive fine pitch variations in speech stimuli) and emotional processing (how well is a given individual able to attribute these variations to a given emotional evaluation) and thus require sophisticated post-hoc controls of the predictive value of a series of acoustical parameters, for example (Lima and Castro, 2011). Using DAVID, it is possible to evaluate sensory accuracy on each individual effect (e.g., pitch shifts ranging from 1 to 100 cents) independently of emotional evaluation, and thus examine individual or group differences on the first sensory stages of emotional processing.
5. Cross-cultural perception: Studies of cross-cultural recognition of emotional vocalizations typically utilize stimuli produced by native speakers of one language, decoded by native speakers of other languages (Scherer et al, 2011), and predict an in-group advantage to decoding expressions produced in your own language compared to others. That advantage can stem either from differences in production (one does not do happiness in the same way in French and Japanese), or to a generic facilitation of emotional processing in semantically-meaningful stimuli. Using DAVID, one can produce cross-cultural stimuli that utilize exactly the same acoustic cues, with the same acoustical strength, and thus control for any possible cultural differences in production.
6. Prosody and comprehension: An ongoing debate in theories of speech comprehension concerns whether lexical analysis occurs independently or conjointly with the processing of non-linguistic, prosodic information (e.g. Halle, 1985; Goldinger, 1998). One experimental issue in investigating this interaction is how to selectively manipulate both aspects of speech production without demand effect on the speakers themselves. Using DAVID, on can create experimental dialog situations where the emotional tone of speech of one or the other speaker is systematically manipulated across two different interpretations of the same utterance (e.g., by using homophones as in Halberstadt et al, 1995) to test whether their dialog partner uses such para-linguistic aspects in addition to linguistic content in order to understand speech.

### 5.2 Music

7. Vocal mimicry: One hypothesis to explain the induction of emotions by instrumental music is that certain acoustic characteristics of music may somehow mimic vocal prosodic expression (Juslin and Västfjäll, 2008). Tests of this hypothesis to date have relied on carefully selected pieces of the classical repertoire judged by the experimenter as good examples of such resemblance (e.g., Vivaldi concerto for two violins). Using DAVID, one could apply acoustical transformations, known to result in emotional expression on voice, to musical instrument samples, and systematically test whether they are perceived with the same emotional characteristics.
8. Singing voice: Emotional expression in opera or pop music singers is an important element of music appreciation (Scherer, 1995)^2^, yet it needs to obey two contradictory constraints: first, the positive relationship between e.g. increased pitch and valence in speech expression; second, the negative relationship between dissonance and valence in music polyphony. By using DAVID on multi-track music recordings, one can generate vocal singing tracks with speechlike emotional transformations while keeping the musical background constant, thus studying this interaction.

#### 5.2.1 Social psychology

9. Attention to social cues: Eye contact and gaze following have been shown to vary as a function of a speaker’s social status and emotional state (e.g. Cui et al, 2014; Gallup et al, 2014). However, these are often confounded in a typical experiment. If a listener looks longer at an angry face, for example, it is unclear if they are doing so because of the emotional state of the speaker per se, or because there is particular visual information in the speaker’s face that they are trying to perceive. Using DAVID, experimenters can set up a video conference between one or more people, and vary the emotional tone of some speaker’s voice in real time, while keeping their visual appearance constant. Listener’s eye movements would reveal whether their attention is being deployed according to emotional auditory cues, or visual facial cues.
10. Emotional stereotypes: One hypothesis that explains the quick and automatic activation of stereotypes based on race, gender, or age, is based on ‘self-fulfilling prophecies’ (Bargh et al, 1996) : if a member of group A holds the stereotype that members of group B are hostile, the former may behave in a more guarded or aggressive manner from the start of an interaction, leading the latter to match this behavior and unwillingly confirm their stereotype. The evidence for this mechanism is only correlational, relying on careful and laborious coding of body language and language use during real social interactions (e.g. Neuberg, 1989; Snyder et al, 1977). With DAVID, one can study causal relationships in such interactions by letting participants A and B talk on the phone after reading a brief description of each other, shifting A and B’s voices in a direction congruent or incongruent with the description, and test whether prejudices persist after the interaction.
11. The Looking glass self: Tice (1992) showed that when people are asked to display a certain emotion before rating their ‘true selves’ on these attributes, their ratings depend on whether or not they thought they were being observed. In other words, people adjust their view of themselves according to how they believe they appear to others. However, With DAVID, one can avoid a demand effect in a similar paradigm where participants are told to either talk to a computer or to a participant in another room. In this case participants need not be explicitly instructed to behave in a certain way.
12. Willingness to cooperate: Van Doorn et al (2012) showed that when participants read a text description of their hypothetical study partners described as happy, participants evaluated their partners as more cooperative when compared to descriptions of angry study partners. It remains unclear whether this effect applies in more realistic ongoing social interactions. Using DAVID, one can create such realistic social interactions and manipulate the expressed emotion at the same time. For example, one can let two participants interact before submitting them to a prisoner’s dilemma paradigm and test how the emotional interaction beforehand influences the participants’ behavior during the social decision-making game
13. Group productivity: In one case study, Parker and Hackett (2012) have described how emotional processes within a group of scientists can influence productivity and creativity. Such studies commonly rely on many hours of data collection and observation. To further study the dynamics and the impact of emotional processes on group productivity in a more controlled fashion, one can use DAVID to manipulate the emotional congruence within group interactions, for instance during a brainstorm session, and analyze how this affects group productivity.

### 5.3 Emotion regulation

14. Expressive writing: Pennebaker and colleagues have found that expressive language use can have beneficial outcomes: people who write about their relationships are more likely to remain in those relationships (Slatcher and Pennebaker, 2006); people who write about their job loss are more likely to be reemployed (Spera et al, 1994) and writing about traumatic experiences improves the health of maximum security inmates (Richards et al, 2000). With DAVID, experimenters could ask participants to read out loud their expressive writing while manipulating the emotional tone of their voice - isolating the role of emotional re-engagement from the effects of factual recollection, language production and other aspects of therapeutic writing.
15. Recollection of past events: Siegman and Boyle (1993) reported high levels of subjective affective and cardiovascular (CV) in participants speaking about emotional events in a mood-congruent voice style. In contrast, when participants spoke with a mood-incongruent voice style, such high levels of CV arousal were not measured. Using DAVID to manipulate the emotional congruence of the voice, one can investigate if participants will have similar lowered affective and cardiovascular arousal when speaking about a sad or fearful event with mood-incongruent speech, as they do when talking about these events with an internally generated emotional voice style, as in Siegman and Boyle (1993).
16. Emotional display: A distinction is often drawn between those emotional expressions of the voice stemming from physiological factors and those that are consciously induced by the speaker to purposely convey an emotion (see e.g. Scherer et al, 2003). In Aucouturier et al (2016), we did not find a feedback effect using the "afraid" manipulation, possibly because its constituent vibrato is not a socially conventional signal (such as raised or lowered f0), but one stemming from a physiological state too distinct from the state the participant is actually in. Using DAVID, it is possible to manipulate vocal properties independently from a participant’s physiological state, and investigate the socially conventional vs. physiological aspects of the vocal expressions of emotion.
17. Mood induction: Mood induction paradigms typically use images of faces (Schneider et al, 1994), cartoons (Falkenberg et al, 2012), or musical extracts (Koelsch et al, 2006). With DAVID it would be possible to develop novel mood induction protocols using the participant’s own voice. This may have additional benefits as the stimuli used are directly related to the participant. In patients with major depression this approach may be of special interest because self-referential processing, which is often altered in depressed patients (see e.g. Segal et al, 1995; Grimm et al, 2009), can be addressed as well.

### 5.4 Peripheral emotional feedback

18. Awareness of the manipulation: In the emotional vocal feedback study by Aucouturier et al (2016), participants were not informed about the manipulation prior to the experiment. Their results showed that, while participants did not detect the manipulation, the emotional transformations done with DAVID were able to induce congruent changes in the participants’ mood. While clinically interesting, it remains an open question whether similar mood changes would occur if participants were made aware of the manipulation, for instance in the context of an intervention. Using DAVID, one could for instance conceive a feedback situation with transformations of a stronger, more noticeable intensity, or where participants receive deliberate instructions that their voice is going to be manipulated.
19. Vocal compensation: In studies where e.g. F0 or F1 feedback is altered (e.g. Houde and Jordan, 1998; Purcell and Munhall, 2006), the general result reported is that participants unknowingly compensate for the perturbations. But such compensation is not always observed (see e.g. Burnett et al, 1998; Behroozmand et al, 2012), and it has been suggested that it may be limited to constrained circumstances, using vowels or monosyllabic words. In Aucouturier et al (2016) no compensations were found for the manipulations, which took place in the context of continuous and more complex speech production. Using DAVID, it is possible to further explore these discrepancies, for instance testing under which circumstances (e.g., different types of social interactions versus self-focused out-loud reading) people may or may not adapt their speech in response to perturbed vocal feedback.
20. Emotional monitoring: In an inferential view of speech production and emotional self-monitoring, it is assumed that voice manipulations will be accepted as self-produced only if there is a sufficient contextual plausibility to the manipulation (e.g. Lind et al, 2014; Aucouturier et al, 2016). DAVID can be used to explore the contextual requirements for accepting the manipulations, informing us about which emotional cues are internalized based on the voice alone and which form part of a larger inferential inferential cluster of cues. For example, it might be that a happy voice transformation is more easily accepted in a friendly social interaction than in an extremely hostile one.

### 5.5 Decision making

21. Moral decision-making: Moral judgments are often influenced by emotional processes. For example, Xiao et al (2015) showed that participants were more sympathetic in their responses to moral dilemmas when they experienced physical pain. Whether the same applies to emotions is an open question. Using DAVID, one could alter the emotional tone of participants’ voices while they read pairs of moral vignettes out loud and let them decide which one is morally worse (e.g. "you see a professor giving a bad grade to a student just because he dislikes him" vs "you see a teenager at a cafeteria forcing a younger student to pay for her lunch", from Clifford et al, 2015). If one vignette is read with a sad voice and the other with a happy or neutral voice, the prediction would be that participants find the vignette that they have read with a negative sounding voice to be the most morally reprehensible.
22. Somatic marker: Building on Damasio’s somatic marker hypothesis (Damasio, 1994), Arminjon and colleagues (2015) asked participants to recall a memorized text while holding a pen between their teeth, finding that such unconscious smiling altered the participants’ emotional ratings of the text but not its episodic memory. One issue here is that, while the smile is not generated explicitely, it is nevertheless initiated actively by the participants. With DAVID, one can let participants recall a memorized test out loud while their voice is made to sound more or less positive, and test whether voice functions as a somatic marker during memory reactivation, without any experimental demand on the participant.
23. Attitudes towards topics: Using false heart-rate feedback, Shahidi and Baluch (1991) showed that in stressful events, participants reported higher levels of embarrassment when they received faster heart-rate feedback. Using DAVID, it would be possible to investigate if hearing oneself with a sad or afraid voice introduces a bias to be less confident in one’s own behavior for example when answering questions on a query: e.g., "this is my answer, but, seeing how I sounded when I said it, I don’t think it’s correct". Similarly, DAVID can be used to investigate if participants become more or less interested in, or more positively or negatively oriented towards a topic of discussion, or to a topic they are reading about (such as a relatively neutral newspaper article), by having them reading it aloud with neutral versus sad or happy voice.

#### 5.5.1 Consumer behaviour

24. Video games: Player satisfaction with online first-person video games is closely related to their ability to let people lose themselves in the game, a phenomenon often referred to as immersion (Jennett et al, 2008; Ryan et al, 2006). Developers of video games use different strategies to enhance game immersion, such as letting several players interact via voice while they are playing, or using music and visual elements to make the game more realistic. Using DAVID, it would be possible to change the voices of interacting players to make the game even more engaging. For example, in a collaborative horror-themed video game, it would be possible to make the players’ voices sound more scared while they are playing, and later measure the effect of the manipulation on the players’ perceived degree of immersion.
25. Karaoke: Vocal processing, such as automatic generation of a harmony voice to second the participant’s singing, is an important focus of the karaoke industry to improve customer enjoyment (Kato, 2000). Using DAVID on singing voice, it is possible to test whether adding realistic emotional expressions in real-time, possibly suited to the target song (e.g. a trembling transformation to a sad song) can enhance the singer’s (and his/her audience’s) appreciation of a performance.

## 5.6 Acknowledgements

This research was funded by a European Research Council Grant StG-335536 CREAM to J.-J.A.

## Footnotes

1 "Da Amazing Voice Inflection Device"' DAVID was so named after Talking Heads' frontman David Byrne' whom we were privileged to count as one of our early users in March 2015.

2 see also http://www.wsj.com/articles/SB10001424052970203646004577213010291701378

## References

Arminjon M, Preissmann D, Chmetz F, Duraku A, Ansermet F, Magistretti PJ (2015) Embodied memory: unconscious smiling modulates emotional evaluation of episodic memories. Frontiers in Psychology 6:650

Astrinaki M, D’alessandro N, Picart B, Drugman T, Dutoit T (2012) Reactive and continuous control ofhmm-based speech synthesis. In: Spoken Language Technology Workshop (SLT), 2012 IEEE, IEEE, pp 252–257

Aucouturier J, Johansson P, Hall L, Mercadié L, Watanabe K (2016) Covert digital manipulation of vocal emotion alter speakers’ emotional states in a congruent direction. Proceedings of the National Academy of Sciences doi:10.1073/pnas.1506552113

Bachorowski JA, Owren MJ (1995) Vocal expression of emotion: Acoustic properties of speech are associated with emotional intensity and context. Psychological science 6(4):219–224

Badian M, Appel E, Palm D, Rupp W, Sittig W, Taeuber K (1979) Standardized mental stress in healthy volunteers induced by delayed auditory feedback (daf). European Journal of Clinical Pharmacology 16(3):171–176

Banziger T, Scherer KR (2006) The role of intonation in emotional expressions. Speech Communication 46:252–267

Bargh JA, Chen M, Burrows L (1996) Automaticity of social behavior: Direct effects of trait construct and stereotype activation on action. Journal of personality and social psychology 71(2):230

Behroozmand R, Korzyukov O, Sattler L, Larson CR (2012) Opposing and following vocal responses to pitch-shifted auditory feedback: Evidence for different mechanisms of voice pitch control. The Journal of the Acoustical Society of America 132(4):2468–2477

Belin P, Fillion-Bilodeau S, Gosselin F (2008) The montreal affective voices: a validated set of nonverbal affect bursts for research on auditory affective processing. Behavior research methods 40(2):531–539

Bertini G, Fontana F, Gonzalez D, Grassi L, Magrini M (2005) Voice transformation algorithms with real time dsp rapid prototyping tools. In: Proceedings of 13th European Signal Processing Conference, Antalya, Turkey

Bestelmeyer PEG, Latinus M, Bruckert L, Rouger J, Crabbe F, Belin P (2012) Implicitly perceived vocal attractiveness modulates prefrontal cortex activity. Cerebral Cortex 22:1263–1270

Boersma P, Weenink D (1996) Praat: doing phonetics by computer (version 5.1.05) [computer program]. Retrieved Nov. 1, 2009, from http://www.praat.org/

Bulut M, Narayanan SS (2008) F0 modifications in emotional speech. J Acoust Soc Am 123(6):4547–4558

Bulut M, Busso C, Yildirim S, Kazemzadeh A, Lee CM, Lee S, Narayanan S (2005) Investigating the role of phoneme-level modifications in emotional speech resynthesis. In: Proceedings of 6th Annual Conference of the International Speech Communication Association (Interspeech), Lisbon, Portugal

Burnett TA, Freedland MB, Larson CR, Hain TC (1998) Voice f0 responses to manipulations in pitch feedback. The Journal of the Acoustical Society of America 103(6):3153–3161

Cabral JP, Oliveira LC (2005) Pitch-synchronous time-scaling for prosodic and voice quality transformations. In: Proceedings of 6th Annual Conference of the International Speech Communication Association (Interspeech), Lisbon, Portugal

Camacho A, Harris JG (2008) A sawtooth waveform inspired pitch estimator for speech and music. The Journal of the Acoustical Society of America 124(3):1638–1652

Clifford S, Iyengar V, Cabeza R, Sinnott-Armstrong W (2015) Moral foundations vignettes: a standardized stimulus database of scenarios based on moral foundations theory. Behavior research methods pp 1–21

Cui G, Zhang S, Geng H (2014) The impact of perceived social power and dangerous context on social attention. PloS one 9(12):e114,077

Damasio AR (1994) Descartes’ error and the future of human life. Scientific American 271(4):144

Eide E, Aaron A, Bakis R, Hamza W, Picheny M, Pitrelli J (2004) A corpus-based approach to <ahem> expressive speech synthesis. In: Proceedings of 5th ISCA Speech Synthesis Workshop, Pittsburg, PA, USA

Elfenbein H, Ambady N (2002) On the universality and cultural specificity of emotion recognition: A meta-analysis. Psychological Bulletin 128(2):203–235

Ethofer T, Anders S, Wiethoff S, Erb M, Herbert C, Saur R, Grodd W, Wildgruber D (2006) Effects of prosodic emotional intensity on activation of associative auditory cortex. Neuroreport 17(3):249–253

Falkenberg I, Kohn N, Schoepker R, Habel U (2012) Mood induction in depressive patients: a comparative multidimensional approach. PloS one 7(1):e30,016-e30,016

Farner S, Veaux c, Beller G, Rodet X, Ach L (2008) Voice transformation and speech synthesis for video games. In: Proceedings of Paris Game Developers Conference, Paris, France

Fleming SM, Lau HC (2014) How to measure metacognition. Frontiers in human neuroscience 8

Gallup AC, Chong A, Kacelnik A, Krebs JR, Couzin ID (2014) The influence of emotional facial expressions on gaze-following in grouped and solitary pedestrians. Scientific reports 4

Godoy E, Rosec O, Chonavel T (2009) Alleviating the one-to-many mapping problem in voice conversion with context-dependent modeling. In: Proceedings of 10th Annual Conference of the International Speech Communication Association (Interspeech), Brighton, UK

Goeleven E, De Raedt R, Leyman L, Verschuere B (2008) The karolinska directed emotional faces: a validation study. Cognition and Emotion 22(6):1094–1118

Goldinger SD (1998) Echoes of echoes? an episodic theory of lexical access. Psychological review 105(2):251

Grimm S, Ernst J, Boesiger P, Schuepbach D, Hell D, Boeker H, Northoff G (2009) Increased self-focus in major depressive disorder is related to neural abnormalities in subcortical-cortical midline structures. Human brain mapping 30(8):2617–2627

Halberstadt JB, Niedenthal PM, Kushner J (1995) Resolution of lexical ambiguity by emotional state. Psychological Science pp 278–282

Halle M (1985) Speculations about the representation of words in memory. Phonetic linguistics pp 101–114

Hammerschmidt K, Jurgens U (2007) Acoustical correlates of affective prosody. Journal of Voice 21(5):531–540

Houde JF, Jordan MI (1998) Sensorimotor adaptation in speech production. Science 279(5354):1213–1216

Inanoglu Z, Young S (2007) A system for transforming the emotion in speech: combining data-driven conversion techniques for prosody and voice quality. In: INTERSPEECH, pp 490–493

Jennett C, Cox AL, Cairns P, Dhoparee S, Epps A, Tijs T, Walton A (2008) Measuring and defining the experience of immersion in games. International journal of human-computer studies 66(9):641–661

Juslin PN, Laukka P (2003) Communication of emotions in vocal expression and music performance: different channels, same code? Psychological bulletin 129(5):770–814

Juslin PN, Västfjäll D (2008) Emotional responses to music: the need to consider underlying mechanisms. The Behavioral and brain sciences 31(5):559–75; discussion 575-621

Juslin PN, Scherer KR, Harrigan J, Rosenthal R, Scherer K (2005) Vocal expression of affect. The new handbook of methods in nonverbal behavior research pp 65–135

Kato H (2000) Karaoke apparatus selectively providing harmony voice to duet singing voices. US Patent 6,121,531

Koelsch S, Fritz T, Müller K, Friederici AD, et al (2006) Investigating emotion with music: an fmri study. Human brain mapping 27(3):239–250

Laukka P, Juslin P, Bresin R (2005) A dimensional approach to vocal expression of emotion. Cognition & Emotion 19(5):633–653

Lima CF, Castro SL (2011) Speaking to the trained ear: musical expertise enhances the recognition of emotions in speech prosody. Emotion 11(5):1021

Lind A, Hall L, Breidegard B, Balkenius C, Johansson P (2014) Auditory feedback of one’s own voice is used for high-level semantic monitoring: the "self-comprehension" hypothesis. Frontiers in human neuroscience 8

MacMillan K, Droettboom M, Fujinaga I (2001) Audio latency measurements of desktop operating systems. In: Proc. of International Computer Music Conference

Mangini MC, Biederman I (2004) Making the ineffable explicit: Estimating the information employed for face classifications. Cognitive Science 28(2):209–226

Boula de Mareüil P, Celerier P, Toen J (2002) Generation of emotions by a morphing technique in english, french and spanish. In: Proceedings of Speech Prosody, Aix-en-Provence, France, pp 187–190

Mayor O, Bonada J, Janer J (2009) Kaleivoicecope: Voice transformation from interactive installations to video-games. In: Proceedings of AES 35th International Conference, London, UK

Moulines E, Charpentier F (1990) Pitch-synchronous waveform processing techniques for text to speech synthesis using diphones. Speech Communications 9:453476

Neuberg SL (1989) The goal of forming accurate impressions during social interactions: attenuating the impact of negative expectancies. Journal of Personality and Social Psychology 56(3):374

Oudeyer PY (2003) The production and recognition of emotions in speech: features and algorithms. International Journal in Human-Computer Studies 59(1-2):157–183

Paquette S, Peretz I, Belin P (2013) The "musical emotional bursts": a validated set of musical affect bursts to investigate auditory affective processing. Frontiers in psychology 4

Parker JN, Hackett EJ (2012) Hot spots and hot moments in scientific collaborations and social movements. American Sociological Review 77(1):21–44

Peirce JW (2007) Psychopy—psychophysics software in python. Journal of neuroscience methods 162(1):8–13

Pittam J, Gallois C, Callan V (1990) The long-term spectrum and perceived emotion. Speech Communication 9:177–187

Purcell DW, Munhall KG (2006) Adaptive control of vowel formant frequency: Evidence from real-time formant manipulation. The Journal of the Acoustical Society of America 120(2):966–977

Richards JM, Beal WE, Seagal JD, Pennebaker JW (2000) Effects of disclosure of traumatic events on illness behavior among psychiatric prison inmates. Journal of Abnormal Psychology 109(1):156

Roebel A (2010) Shape-invariant speech transformation with the phase vocoder. In: INTERSPEECH, pp 2146–2149

Roesch EB, Tamarit L, Reveret L, Grandjean D, Sander D, Scherer KR (2011) Facsgen: a tool to synthesize emotional facial expressions through systematic manipulation of facial action units. Journal of Nonverbal Behavior 35(1):1–16

Rosenthal R, Rubin DB (1989) Effect Size Estimation for One-Sample Multiple-Choice-Type Data : Design, Analysis, and Meta-Analysis. Psychological Bulletin 106(2):332–337

Russ JB, Gur RC, Bilker WB (2008) Validation of affective and neutral sentence content for prosodic testing. Behavior research methods 40(4):935–939

Ryan RM, Rigby CS, Przybylski A (2006) The motivational pull of video games: A self-determination theory approach. Motivation and emotion 30(4):344–360

Sato W, Kochiyama T, Yoshikawa S, Naito E, Matsumura M (2004) Enhanced neural activity in response to dynamic facial expressions of emotion: an fmri study. Cognitive Brain Research 20(1):81–91

Scherer K (2003) Vocal communication of emotion: A review of research paradigms. Speech Communication 40(1-2):227–256

Scherer K, Oshinsky J (1977) Cue utilization in emotion attribution from auditory stimuli. Motivation and Emotion 1:331–346

Scherer KR (1987) Vocal assessment of affective disorders. In: Maser JD (ed) Depression and expressive behavior, Hillsdale, New Jersey: Erlbaum, pp 57–82

Scherer KR (1995) Expression of emotion in voice and music. Journal of voice 9(3):235–248

Scherer KR, Johnstone T, Klasmeyer G (2003) Vocal expression of emotion. Handbook of affective sciences pp 433–456

Scherer KR, Clark-Polner E, Mortillaro M (2011) In the eye of the beholder? universality and cultural specificity in the expression and perception of emotion. International Journal of Psychology 46(6):401–435

Schneider F, Gur RC, Gur RE, Muenz LR (1994) Standardized mood induction with happy and sad facial expressions. Psychiatry research 51(1):19–31

Segal ZV, Gemar M, Truchon C, Guirguis M, Horowitz LM (1995) A priming methodology for studying self-representation in major depressive disorder. Journal of Abnormal Psychology 104(1):205

Shahidi S, Baluch B (1991) False heart-rate feedback, social anxiety and self-attribution of embarrassment. Psychological reports 69(3):1024–1026

Siegman AW, Boyle S (1993) Voices of fear and anxiety and sadness and depression: the effects of speech rate and loudness on fear and anxiety and sadness and depression. Journal of Abnormal Psychology 102(3):430

Slatcher RB, Pennebaker JW (2006) How do i love thee? let me count the words the social effects of expressive writing. Psychological Science 17(8):660–664

Snyder M, Tanke ED, Berscheid E (1977) Social perception and interpersonal behavior: On the self-fulfilling nature of social stereotypes. Journal of Personality and Social Psychology 35(9):656

Spera SP, Buhrfeind ED, Pennebaker JW (1994) Expressive writing and coping with job loss. Academy ofMan-agement Journal 37(3):722–733

Tice DM (1992) Self-concept change and self-presentation: the looking glass self is also a magnifying glass. Journal of personality and social psychology 63(3):435

Toda T, Muramatsu T, Banno H (2012) Implementation of computationally efficient real-time voice conversion. In: INTERSPEECH, Citeseer

Todorov A, Dotsch R, Porter JM, Oosterhof NN, Falvello VB (2013) Validation of data-driven computational models of social perception of faces. Emotion 13(4):724

Van Doorn EA, Heerdink MW, Van Kleef GA (2012) Emotion and the construal of social situations: Inferences of cooperation versus competition from expressions of anger, happiness, and disappointment. Cognition & emotion 26(3):442–461

Wagner HL (1993) On measuring performance in category judgment studies of nonverbal behavior. Journal of Nonverbal Behavior 17(1):3–28

Xiao Q, Zhu Y, Luo Wb (2015) Experiencing physical pain leads to more sympathetic moral judgments. PloS one 10(10):e0140,580

